# Arabidopsis RabGDIs are essential for the asymmetric division of zygotes and embryonic patterning

**DOI:** 10.1101/2025.06.23.660972

**Authors:** Gui-Min Yin, Ya-Nan Wu, Weiqi Wang, Shan-Shan Dun, Sha Li, Liwen Jiang, Zi-zhen Liang, Yan Zhang, Feng Xiong

## Abstract

An asymmetric division of zygotes sets up the apical-basal body axis and starts the life cycle of angiosperms. In addition to specific expression of transcription factors and polar distribution of auxin, vesicular trafficking-mediated endomembrane dynamics are critical for zygotic division and embryonic patterning. As key regulators of vesicular trafficking, whether Rab GTPases play a role in these processes is unclear. We report that Arabidopsis guanine nucleotide dissociation inhibitors for Rab (RabGDIs) are essential for the asymmetric division of zygotes and embryonic patterning likely through regulating the dynamic targeting of canonical Rab GTPases, especially Rab5. Arabidopsis RabGDIs specifically interact with canonical Rab GTPases. Functional loss of Arabidopsis *RabGDI1* and *RabGDI2* compromises the asymmetry of zygotic division, cell fate determination, and embryonic patterning. Polar distribution of vacuolar dynamics was disrupted in zygotes and 1-cell embryo of *gdi1 gdi2*, suggesting a key role of RabGDIs in vacuolar dynamics. Furthermore, genetic interference of canonical Rab5, a key regulator of vacuolar trafficking and biogenesis, in zygotes leads to similar defects, consistent with the disturbed localization of Rab5 in *gdi1 gdi2* embryos. Results presented demonstrate the key role of RabGDIs through Rab GTPases in asymmetric division of zygotes and embryonic patterning.

## Introduction

In angiosperms, a sperm and an egg fuse to form a zygote, which undergoes a series of oriented and highly ordered mitotic divisions and differentiation to form a multicellular embryo. The first zygotic division is asymmetric, which leads to a small apical cell and a large basal cell (Mansfield & Briarty, 1991; Ueda & Laux, 2012; Dresselhaus & Jurgens, 2021). The apical cell will give rise to the embryo whereas the basal cell forms extra-embryonic suspensor, which later undergoes degeneration and contributes to the embryonic root formation (Lau *et al*., 2012; Dresselhaus & Jurgens, 2021). The establishment of apical-basal polarity and tissue-specific differentiation during early embryogenesis requires intricate coordination between transcriptional regulators (Haecker *et al*., 2004; Breuninger *et al*., 2008; Zhang *et al*., 2017) and hormone gradients (Dhonukshe *et al*., 2008; Liu *et al*., 2015; Robert *et al*., 2018). Differential expression of transcription factors such as WOXs, as well as polar distribution of auxin in the apical or basal cell of 1-cell embryo are essential for following divisions, which lead to proper patterning of embryos (Friml *et al*., 2003; Haecker *et al*., 2004; Breuninger *et al*., 2008; Rademacher *et al*., 2012; Robert *et al*., 2013; Zhang *et al*., 2017; Robert *et al*., 2018).

Vesicular trafficking-mediated endomembrane dynamics are critical for zygotic division and embryonic patterning (Dhonukshe *et al*., 2008; Jiang *et al*., 2020; Matsumoto *et al*., 2021). Anterograde trafficking is hinted as a key factor for zygotic division and embryonic patterning. Polar distribution of auxin in 1-cell embryo as well as during embryonic patterning depends on asymmetric distribution of PIN-formed (PINs) at the plasma membrane (PM) (Friml *et al*., 2003), which requires post-Golgi secretion (Geldner *et al*., 2003; Kleine-Vehn & Friml, 2008). In addition to exocytic trafficking, vacuolar dynamics play a key role in zygotic division and embryonic patterning. Right after fertilization, vacuoles in zygotes form tubular strands around the nucleus and gradually fuse into a large vacuole at the base, which in turn drives the nucleus upward (Kimata *et al*., 2019; Matsumoto *et al*., 2021). The positioning of vacuoles and the nucleus is crucial for the asymmetric division of zygotes (Jiang *et al*., 2020; Kimata & Ueda, 2020; Matsumoto *et al*., 2021). Arabidopsis mutants with defective vacuolar dynamics often results in defective embryogenesis (Kimata *et al*., 2019; Jiang *et al*., 2020; Matsumoto *et al*., 2021).

Rab GTPases are key regulators of vesicular trafficking among endomembrane compartments (Nielsen *et al*., 2008). There are at least 57 members of Rab GTPases in *Arabidopsis*, grouped into eight subfamilies (Vernoud *et al*., 2003), participating in multiple developmental processes and stress responses (Tripathy *et al*., 2021; Lu *et al*., 2024). Most Rab GTPases are canonical, i.e. modified at the C-termini by geranylgeranyl anchors and conserved among different phyla (Tripathy *et al*., 2021). There are, however, plant-unique Rab GTPases, such as Arabidopsis ARA6, that is modified by *N*-myristoylation and *S*-acylation at the N-terminus (Ueda *et al*., 2001). Rab GTPases cycle between GTP-bound active and GDP-bound inactive status, a process controlled by guanine nucleotide exchange factors (GEFs) and GTPase activating proteins (GAPs) (Muller & Goody, 2018).

In order to mediate vesicular trafficking between two compartments, Rab GTPases are dynamically trafficked between donor/resident compartments and target compartments. Yeast and metazoan studies indicate lipid-anchored Rab GTPases are escorted to their donor/resident membranes by Rab Escort Proteins (REPs) and from there, Rab GTPases mediate vesicular trafficking to their target membrane (Leung *et al*., 2007; Gutkowska *et al*., 2015; Shinde & Maddika, 2018). After fulfilling the roles in vesicle delivery, Rab GTPases are extracted by guanine nucleotide dissociation inhibitors (RabGDIs) from their target membrane, stabilized in cytoplasm, and recycled to donor/resident membrane (Alory & Balch, 2003; Seabra & Wasmeier, 2004; Muller & Goody, 2018; Nielsen, 2020). Although RabGDIs have been identified in plants (Zarsky *et al*., 1997; Ueda *et al*., 1998; Liu *et al*., 2020) and Arabidopsis RabGDI1/2 complement the yeast *gdi1* mutant (Zarsky *et al*., 1997; Ueda *et al*., 1998), whether and how RabGDIs mediate the targeting of Rab GTPases are unclear.

We report here that Arabidopsis RabGDIs interact with canonical Rab GTPases. Functional loss of two constitutively expressed members, *GDI1* and *GDI2*, results in embryo lethality. The first zygotic division is compromised in *gdi1 gdi2* such that zygotic polarity and division asymmetry are affected. We further demonstrate that both polar auxin distribution and specific expression of *WOX8* are impaired in *gdi1 gdi2*, leading to defective embryonic patterning. By genetic interfering with major Rab GTPases, we demonstrate that Rab5-mediated vacuolar trafficking is crucial for zygotic division and embryonic patterning.

## Materials and Methods

### Plant Materials and Growth Conditions

Arabidopsis Columbia-0 ecotype was used as the wild type for all experiments. *DR5:erGFP* (Yin *et al*., 2024b), *PIN1g:GFP* (Yin *et al*., 2024b), *pWOX8gΔ:NLS-vYFP3* (Xiong *et al*., 2019)–marker lines were described. The mutants *gdi1* and *gdi2* were generated by CRISPR/Cas9. Floral dip transformation was mediated by *Agrobacterium* strain GV3101 (WEIDI Biotech, Cat# AC1001) as described (Yin *et al*., 2024a), and stable transgenic plants, *GDI1g:GUS*, *GDI2g:GUS*, *GDI3g:GUS*, *UBQ10:mRFP-GDI1*, *UBQ10:mRFP-GDI2, proGDI1:mRFP-GDI1*, *proGDI2:mRFP-GDI2*, *UBQ10:GFP-ARA7*, *proEC1.1:mRFP-ARA7^DN^*, *proEC1.1:RFP-RabG3f^DN^*, and *proEC1.1:GFP-RabA4b^DN^*were selected on half-strength MS medium (1/2 MS) supplemented with 30 µg/ml Basta salts (Sigma-Aldrich) or 25 µg/ml Hygromycin (Roche). Surface-sterilized seeds with chlorine (produced by NaClO and HCl) for 6 hrs were plated on 1/2 MS medium. Plates were kept at 4°C in darkness for 3 days before being transferred to a growth chamber with a 16-h-light/8-h-dark photoperiod at 22°C. 7-day seedlings were planted in substrate containing nutrient soil and vermiculite (v/v = 1:1) and cultured in the plant artificial climate chamber with constant temperature (22°C), humidity (70%) and photoperiod (16 h light/8 h dark, 200 μmol m^−2^ s^−1^ cool-white fluorescent illumination).

### DNA manipulation

The Gateway system (Invitrogen) was used for plasmid construction unless noted otherwise. The pENTR/D/TOPO vector (Invitrogen) was used to generate entry vectors. Entry vectors of coding sequences (CDS) were generated by using the following primer pairs: ZP8397/ZP9497 for *GDI1*, ZP8399/ZP9498 for *GDI2*, ZP8401/ZP9499 for *GDI3*, ZP927/ZP928 for *RabG3b*, ZP9338/ZP9340 for *ARA4*, PZ1/ZP8378 for *ARA7*, J717/J718 for *ARA7^DN^*, ZP1390/ZP2039 for *ARA6*, ZP3762/ZP3763 for *RabA4b*, ZP1089/ZP1090 for *RabA4b^DN^*, ZP2545/ZP2546 for *RabG3f,* ZP2739/ZP2740 *of RabG3f^DN^* and ZP492/493 for *ROP2*.

GUS reporter constructs were generated through homologous recombination by using pEASY-Uni Seamless Cloning and Assembly Kit (TransGen Biotech, Cat# CU101). Promoter sequences of *RabGDIs* were amplified with the following primer pairs: ZP9867/ZP9868 for *proGDI1*, ZP9871/ZP9872 for *proGDI2*, and ZP9875/ZP9876 for *proGDI3*. Genomic sequences of *RabGDIs* were amplified with the following primers: ZP9869/ZP9870 for *GDI1g*, ZP9873/ZP9874 for *GDI2g*, ZP9877/ZP9878 for *GDI3g*. The resultant PCR fragments were inserted into the destination vector *GW:GUS-gene of interest* using *Bam*HI/*Xba*I and *Sac*I in two-step cloning to generate *proGDI1:GUS-GDI1* (*GDI1g:GUS*), *proGDI2:GUS-GDI2* (*GDI2g:GUS*), and *proGDI3:GUS-GDI3* (*GDI3g:GUS*).

The expression vectors for complementation assays, *UBQ10:mRFP-GDI1*, *UBQ10:mRFP-GDI2, proGDI1:mRFP-GDI1*, and *proGDI2:mRFP-GDI2*, were generated through homologous recombination. The CDS of *RabGDIs* fragments were amplified with the following primers: ZP10584/ZP10585 for *GDI1* and ZP10586/ZP10587 for *GDI2*. The resultant PCR fragments were recombined with *UBQ10:mRFP* (Hao *et al*., 2023) pre-digested with *Sac*I to generate the vectors *UBQ10:mRFP-GDI1* and *UBQ10:mRFP-GDI2,* respectively. The promoter sequences and genomic sequences of *RabGDIs* were amplified with the following primer pairs: P4564/ P4565 for *proGDI1*, P4566/ P4567 for *proGDI2*, P4568/ P4569 for *GDI1g*, P4570/ P4571 for *GDI2g*. PCR fragments of *RabGDIs* promoter sequences and genomic sequences were recombined with *UBQ10:mRFP* (Hao *et al*., 2023) using *Sal*I/*Spe*I and *Bam*HI in two-step cloning to generate *proGDI1:mRFP-GDI1* or *proGDI2:mRFP-GDI2*.

The expression construct *pEC1.1:mRFP-ARA7^DN^*was generated as followed: *pEC1.1* was amplified with the primer pair P5020/P5021; the resultant PCR fragments were recombined with *UBQ10:mRFP* (Hao *et al*., 2023) pre-digested with *Sal*I/*Spe*I to generate *pEC1.1:mRFP*; a *ARA7^DN^*fragment amplified with the primer pair P5026/P5027 was inserted into *pEC1.1:mRFP* pre-digested with *Bam*HI to generate *pEC1.1:mRFP-ARA7^DN^* (Hao *et al*., 2024). For *pEC1.1:GFP-RabG3f^DN^*, *pEC1.1* fragments were inserted into *pCAMBIA1300-GFP-Nos* through *Pst*I/*Bam*HI-double digestion and T4-ligation to generate *pEC1.1:GFP; RabG3f^DN^* fragment amplified with the primer pair PZ201/PZ202 was recombined with *pEC1.1:GFP* pre-digested with *Sac*I to generate *pEC1.1:GFP-RabG3f^DN^* (Cui *et al*., 2014). For *pEC1.1:GFP-RabA4b^DN^*, the *pEC1.1* fragments amplified with the primer pair PZ231/PZ233 were recombined with *35S:GFP-GW* pre-digested with *Hind*III/*Spe*I to generate destination vector *pEC1.1:GFP-GW*; the entry vector for *RabA4b^DN^* was combined with *35S:GFP-GW* using Gateway LR Clonase™ II Enzyme Mix. *UBQ10:GFP-ARA7* were generated by combination using the entry vector for *ARA7* with the destination vector *UBQ10:GFP-GW* (Xiong *et al*., 2019).

To generate the CRISPR/Cas9 construct, *RabGDIs*-specific target sequences were amplified with the primers ZP8219/ZP8220/ZP8221/ZP8222, and *pCBC-DT1T2* was used as the template. Purified PCR fragments were inserted into *pHSE401E* using restriction-ligation reactions as described (Rao *et al*., 2021) to generate the construct *pHSE401E-RabGDI-sgRNA-Cas9*. To verify genomic editing of *GDI1/2*, the primer pair P3705/PZ234 and ZP9249/PZ235 were used to amplify *GDI1* and *GDI2* in CRISPR/Cas9-transformed plants, respectively. The resultant PCR fragments were sequenced with the primer P3705 for *GDI1* and ZP9250 for *GDI2*.

To generate vectors used in 2in1 BiFC assays, CDSs were amplified with the following primer pairs: ZP11516/P216 for *GDI1*, ZP11518/P217 for *GDI2*, ZP11520/P218 for *GDI3*, ZP78Y/ZP79Y for *SCN1/RhoGDI1*, ZP11522/ZP11523 for *ARA7*, P1133/P1134 for *ARA6*, P208/P209 for *ARA4*, P210/P211 for *ROP2*, P212/P213 for *RabG3b*, and ZP28Y/ZP29Y for *RabA4b*. CDSs of *GDI1*, *GDI2*, *GDI3*, and *SCN1/RhoGDI1* were recombined with *pDONR221-P1P4-TOPO* vector while the coding sequences of *ARA7*, *ARA6*, *ARA4*, *ROP2*, *RabG3b*, and *RabA4b* were recombined with *pDONR221-P3P2-TOPO* by BP reactions. Different combinations of entry vectors in *pDONR221-P1P4-TOPO* and *pDONR221-P3P2-TOPO* were used in LR reactions with the destination vector *pBiFCt-2in1-NN* or *pBiFCt-2in1-NC* (Grefen & Blatt, 2012; Pang *et al*., 2022) to generate expression vectors used in the 2in1 BiFC system.

To generate expression vectors used in FRET assays, The entry vectors of *RabGDIs* and different Rab GTPases were combined with the destination vector *35S:GFP-GW* using Gateway LR Clonase™ II Enzyme Mix (Invitrogen, Cat# 11791020) to generate expression vectors *35S:GFP-GDI1*, *35S:GFP-GDI2*, and *35S:GFP-GDI3*, respectively. Entry vectors for *ARA6, ARA7*, *RabG3b*, *ARA4*, *RabA4b,* and *ROP2* were used in LR reactions with the destination vectors *35S:GW-mCherry* or *35S:mCherry-GW* to generate the expression vectors *35S:ARA6-mCherry*, *35S:mCherry-ARA7*, *35S:mCherry-RabG3b*, *35S:mCherry-ARA4*, *35S:mCherry-RabA4b*, and *35S:mCherry***-** *ROP2*.

All PCR amplifications were performed with Phusion hot-start high-fidelity DNA polymerase with the annealing temperature and extension times recommended by the manufacturer (Thermo Fisher Scientific, Cat# F548S). Entry vectors were sequenced, and sequences were analyzed using Vector NTI (https://www.invitrogen.com/VectorNTI) or DNAMAN version 6 (http://www.lynnon.com/dnaman.html). All primers are listed in Supplemental Table 1.

### Quantitative real-time PCRs (RT-qPCRs)

Extraction of total RNAs from different tissues were performed with Ultrapure RNA Kit (CWBIO, Cat# CW0597S). Oligo (dT)-primed cDNAs were synthesized using ReverTra AceTM qPCR RT Master Mix with gDNA Remover Kit (TOYOBO, Cat# FSQ-301). qPCRs were performed with the ABI QuantStudio 6 Flex using PerfectStart Green qPCR SuperMix(+Dye II) (TransGen Biotech, Cat# AQ602). Primers used in RT-qPCRs are ZP9500/ZP9501 for *GDI1*, ZP9502/ZP9503 for *GDI2*, ZP9504/ZP9505 for *GDI3,* and ZP687/ZP688 for *GAPDH* as an internal control. All primers are listed in Supplemental Table 1.

### Histochemical GUS staining

Tissues from GUS reporter plants were stained with X-Gluc (Sigma-Aldrich, Cat# 203783) substrates at 37°C for 2 hrs to 24 hrs, transferred to 70% ethanol to remove the residual color, and imaged by an Olympus BX51 microscope (Olympus, Tokyo, Japan) or an Olympus SZX16 microscope (Olympus, Tokyo, Japan).

### Protein interaction assays

BiFCs were performed in tobacco (*Nicotiana tabacum*) by transient transformations as described (Li *et al*., 2021) and P19 protein was used to suppress gene silencing (Yin *et al*., 2024a). The transformed tobacco plants were cultured for 48-72 hrs before imaging.

FRET assays were performed in Arabidopsis protoplasts by transient transformations as described (Yin *et al*., 2024a). Calculation of FRET efficiency is as described (Li *et al*., 2021). Energy transfer efficiency (E) was calculated according to the formula: E=1-[e/(e+f-eb/a-cg/d)], “eb/a” used to calculate donor spectral bleed-through (DSBT), “cg/d” used to calculate acceptor spectral bleed-through (ASBT). The E values of GFP-fusions/mCherry-fusions with GFP-fusions/mCherry alone were compared to determine positive interactions. Parameters are: a, excitation/emission wavelengths set to 488 nm/505-550 nm for GFP-RabGDIs; b, excitation/emission wavelengths set to 488 nm/575-656 nm for GFP-RabGDIs; c, excitation/emission wavelengths set to 488 nm/575-656 nm for mCherry-fusion; d, excitation/emission wavelengths set to 561 nm/575-656 nm for mCherry-fusion; e, excitation/emission wavelengths set to 488 nm/505-550 nm for GFP-RabGDIs and mCherry-fusion; f, excitation/emission wavelengths set to 488 nm/575-656 nm for GFP-RabGDIs and mCherry-fusion; g, excitation/emission wavelengths set to 561 nm/575-656 nm for GFP-RabGDIs and mCherry-fusion.

### Ovule clearing and Differential interference contrast (DIC) imaging

Ovules dissected from siliques were fixed in 4% Glutaraldehyde in PBS Buffer and cleared with Hoyer’s solution (chloral hydrate: glycerol: water = 8: 1: 2, w: v: v) following the protocol described (Yadegari *et al*., 1994). A Zeiss LSM880 laser scanning microscope with DIC optics was used to capture the images of cleared embryos.

### Vacuole observation using confocal laser scanning microscopy

Ovules after fertilization were fixed in 4% Glutaraldehyde in PBS Buffer at 4°C for 12 hrs after applying three times vacuum, 30 mins each time. Then the samples were washed three times by PBS Buffer and followed by a gradient dehydration in 10%, 20%, 30%, 50%, 70%, 80%, 90% and 95% ethanol for 20 min each step. After immersion in new 95% ethanol for 12 h, the samples were cleared in mixture solution (benzyl benzoate: benzyl alcohol = 2: 1, v: v) for another 12 h. Finally, the zygotes were observed following excitation with laser light (488 nm).

### Fluorescence microscopy

Fluorescence images were captured using a Zeiss LSM880 confocal laser scanning microscope (CLSM). For confocal laser scanning micrograph imaging of GFP, YFP, and mCherry, excitations/emissions are 488 nm/505-550 nm, 514 nm/520-550 nm and 561 nm/575-650 nm, respectively. Double fluorescent labeled materials were captured alternately using line-switching with the multi-track function. DIC imaging of ovules was performed using a Zeiss AxioPhot microscope with DIC optics. Image processing was performed with the Zeiss LSM image processing software (Zeiss).

### Statistical analysis

Quantification data are analyzed by using GraphPad Prism 9 (www.graphpad.com/scientific-software/prism/). All statistical analyses were performed with either Students’ *t*-test or One-Way ANOVA (Tukey’s multiple comparisons test) with build-in analysis tools and parameters. The statistical details, including exact value of n for each measurement, the type of statistical significance test, the definition of significance or P values are all presented in the figure legends or figures.

### Accession Numbers

Arabidopsis Genome Initiative locus identifiers for the genes mentioned in this article are as follows: At2g44100 for *RabGDI1*, At3g59920 for *RabGDI2*, At5g09550 for *RabGDI3*, At4g19640 for *ARA7*, At3g54840 for *ARA6*, At1g22740 for *RabG3b*, At3g18820 for *RabG3f*, At2g43130 for *ARA4*, At4g39990 for *RabA4b*, At1g20090 for *ROP2*, At3g07880 for *SCN1/RhoGDI1*, At1g76750 for *EC1.1*, At1g73590 for *PIN1*, and At5g45980 for *WOX8*.

## Results

### Arabidopsis RabGDIs interact with canonical Rab GTPases

Three RabGDIs are encoded in Arabidopsis genome (Liu *et al*., 2020). RabGDI1 was reported to interact with ARA4 in yeast (Ueda *et al*., 2000). To test whether RabGDIs interacted with Rab GTPases *in planta*, we performed fluorescence resonance energy transfer (FRET) assays. Out of over 57 members of Arabidopsis Rab GTPases, we selected two members of the Rab11 clade for post-Golgi secretion, RabA4b (Preuss *et al*., 2004) and ARA4 (Kirchhelle *et al*., 2016), ARA7 as a canonical Rab5 (Minamino & Ueda, 2019), Rab3Gb as a representative of Rab7 (Cui *et al*., 2014; Minamino & Ueda, 2019). The plant-unique Rab5, ARA6 (Ueda *et al*., 2001), and ROP2, a representative of a different type of small GTPases (Vernoud *et al*., 2003), were used as controls for canonical Rab GTPases. By FRET assays, we determined that all three RabGDIs interacted with all canonical Rab GTPases tested but not ARA6 or ROP2 (Fig. 1a-c). FRET signals between RabGDIs and canonical Rab GTPases were significantly higher than those between RabGDIs and ARA6 or ROP2 (Fig. 1d-f). To provide further evidence supporting the interaction between RabGDIs and canonical Rab GTPases, we performed bimolecular fluorescence complementation (BiFC) assays. Results from BiFC assays further verified the interaction between RabGDIs and canonical Rab GTPases but not the plant-unique ARA6 or ROP2 (Fig. S1). In addition, canonical Rab GTPases did not interact with the GDI of ROPs (Feng *et al*., 2016), RhoGDI1/SCN1 (Fig. S1), supporting the RabGDIs-canonical Rab GTPases interaction specificity.

**Figure 1.**
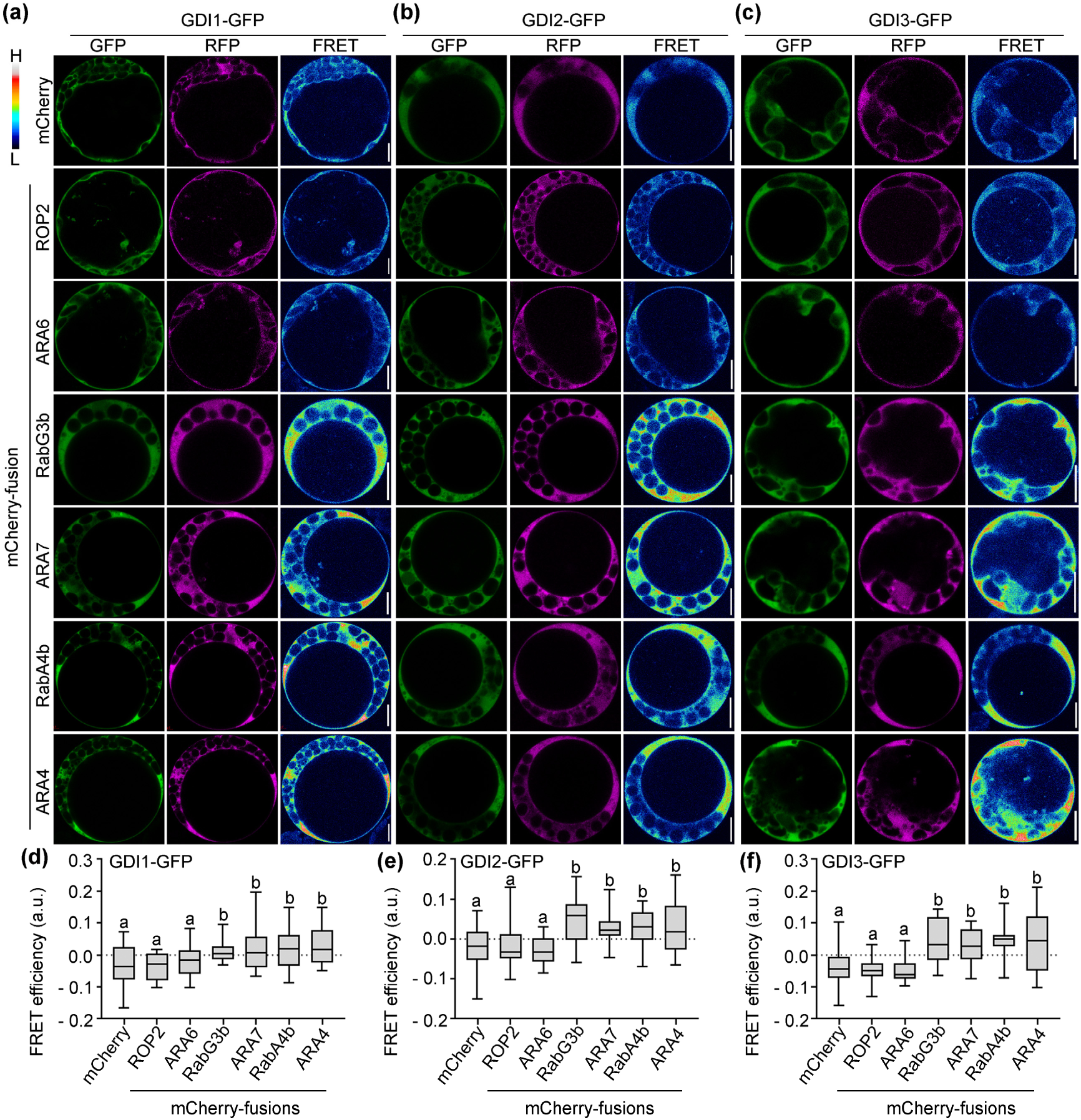
RabGDIs interact with canonical Rab GTPases. Confocal laser scanning microscopic (CLSM) images of FRET assays (a-c) or quantification of FRET efficiency (d-f). FRET signals are represented in pseudo-color, covering the full range of measures values within each dataset (high to low, upper left corner). Results are means ± SD (n > 30). Each combination was examined with three replicate experiments. Different letters indicate significantly different groups (OneWay ANOVA, Tukey’s multiple comparisons test, P<0.01). a.u. stands for arbitrary fluorescence unit. Bars = 10 µm.

### *RabGDI1*/*2* are constitutively expressed

We first examined the expression of the three Arabidopsis *RabGDI*s by reverse transcription quantitative PCRs (RT-qPCRs). *GDI1* and *GDI2* were constitutively expressed in various tissues, such as seedlings, leaves, flowers and primary roots (Fig. S2), as reported (Zarsky *et al*., 1997; Ueda *et al*., 1998). By contrast, *GDI3* was detected only in floral tissues, with a peak expression in mature pollen (Fig. S3). To verify the expression pattern of *GDI1*, *GDI2*, and *GDI3*, we generated genomic-GUS reporter transgenic plants for *GDI1*, *GDI2*, and *GDI3.* By histochemical GUS staining of *GDI1g:GUS*, *GDI2g:GUS*, or *GDI3g:GUS* plants, we determined that *GDI1* and *GDI2* were constitutively expressed in various tissues (Fig. S2 and Fig. S3) whereas *GDI3* was hardly detectable in vegetative tissues (Fig. S3). Overlapping as well as differential expression of three RabGDI members suggests functional redundancy and diversification.

### Functional loss of *GDI1* and *GDI2* results in embryo lethality

To investigate the biological functions of *GDI1* and *GDI2* in *Arabidopsis*, we took a reverse genetic approach by generating *GDI* mutants using CRISPR/Cas-9-mediated genomic editing (Xing *et al*., 2014). We identified *gdi1*, *gdi2*, *gdi1*/+ *gdi2*, and *gdi1 gdi2*/+, from Cas9-edited and T-DNA-free plants (Fig. S4), in which a 1 base-pair deletion resulted in pre-stop codon in both *GDI1* and *GDI2* genomic loci (Fig. 2a-b). No homozygous *gdi1 gdi2* plants were identified and siliques of the *gdi1*/+ *gdi2* or *gdi1 gdi2*/+ plants contained around 25% defective seeds (Fig. 2d-e, 2h), unlike wild type (Fig. 2c, 2h). These results suggested embryo lethality of *gdi1 gdi2*. To test this idea, we examined the segregation of self-fertilized *gdi1*/+ *gdi2* or *gdi1 gdi2*/+ plants. Indeed, no homozygous *gdi1 gdi2* plants were obtained from self-fertilized *gdi1*/+ *gdi2* or *gdi1 gdi2*/+ plants (Table 1), supporting embryo lethality of *gdi1 gdi2*. Functional loss of *GDI1* or *GDI2* alone did not affect seed set (Fig. 2h, Table 1), suggesting a redundancy between *GDI1* and *GDI2*. In addition, *gdi1 gdi2* transmitted comparably to that of wild type either through male or female gametophytes (Table 1), indicating that functional loss of *GDI1* and *GDI2* did not affect gametophytes. To further verify that functional loss of *GDI1* and *GDI2* resulted in embryo lethality, we introduced the genomic mRFP-fusions of *GDI1/2*, *proGDI1:mRFP-GDI1* and *proGDI2:mRFP-GDI2,* into *gdi1*/+ *gdi2* or *gdi1 gdi2*/+ plants. We obtained *proGDI1:mRFP-GDI1 gdi1 gdi2* plants and *proGDI2:mRFP-GDI2 gdi1 gdi2* plants (Fig. S5) in which seed set was comparable to that of wild type (Fig. 2f-h), indicating the full complementation of *gdi1 gdi2* by either *mRFP-GDI1* or *mRFP-GDI2*.

**Figure 2.**
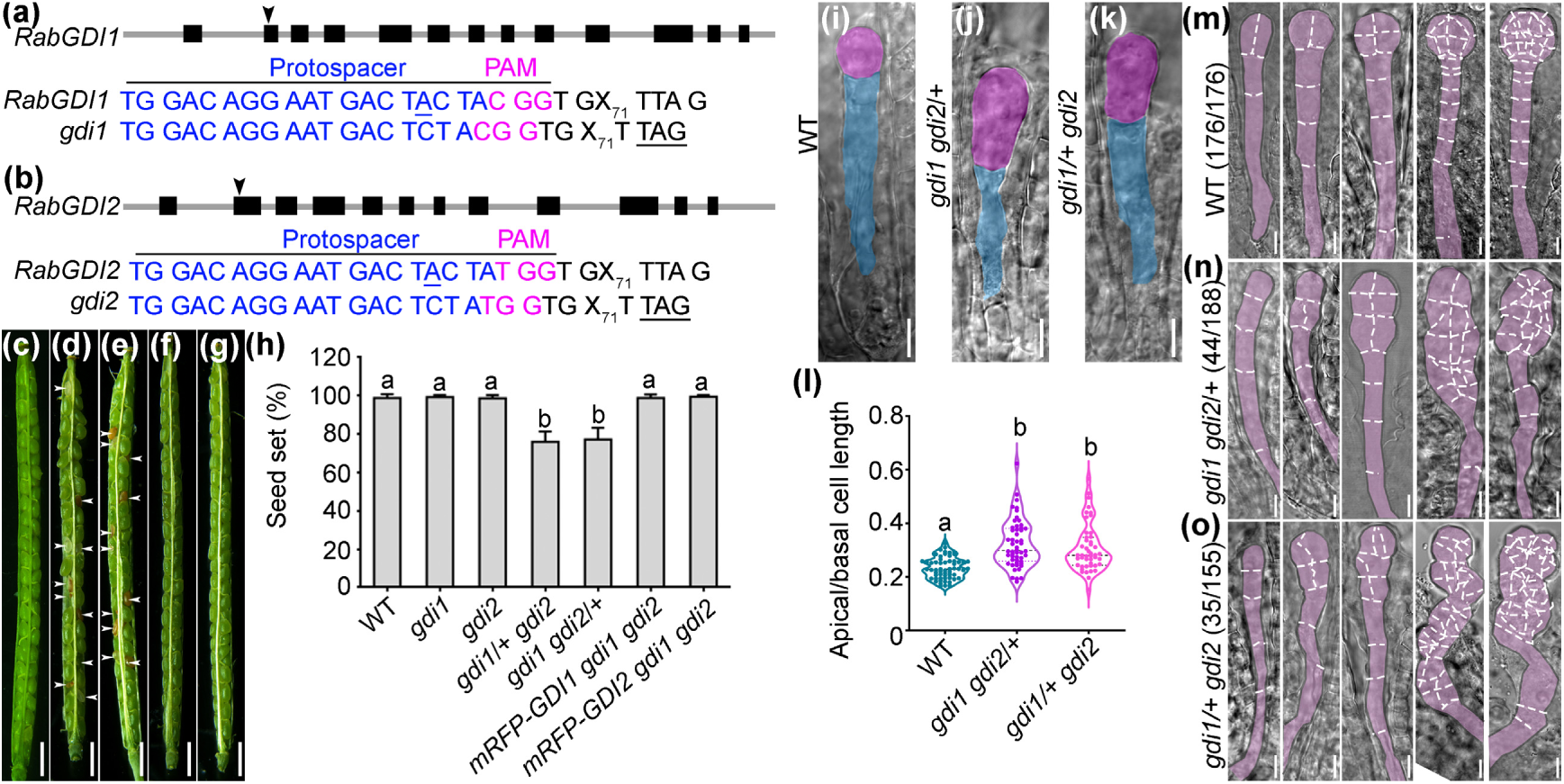
Functional loss of *RabGDI1* and *RabGDI2* results in embryo lethality. (a-b) Genomic structure of *RabGDI1* (a) and *RabGDI2* (b). The target site of Cas9 is indicated by arrowheads on the genomic loci. PAM sequences and protospacer sequences are indicated by magenta and blue letters, respectively. Cas9-generated base pair deletion (blue, underlined) and the pre-stop codon introduced during genomic editing (black, underlined) are shown. X71 indicates 71 base pairs. Note that the same mutation was generated in *RabGDI1* and *RabGDI2* by genome editing. (c-g) Representative siliques from self-pollinated wild type (c), *gdi1*/+ *gdi2* (d), *gdi1 gdi2*/+ (e), *mRFP-RabGDI1 gdi1 gdi2* (f), or *mRFP-RabGDI2 gdi1 gdi2* (g). Arrowheads point at aborted seeds. (h) Seed set. Results are means ± SD (n = 20-30). Different letters indicate significantly different groups (OneWay ANOVA, Tukey’s multiple comparisons test, P<0.01). (i-k) Differential interference contrast (DIC) images of 1-cell embryo from wild type (i), *gdi1 gdi2*/+ (j), or *gdi1*/+ *gdi2* (k). The apical cell and basal cell are pseudo-colored in pink and blue, respectively. (l) Ratio of apical/basal cell length at 1-cell embryos. Results shown are means ± SD (SD, n= 46-58). (m-o) Representative DIC images of embryos from 2-cell stages to globular stages from self-fertilized wild-type (m), *gdi1 gdi2*/+ (n), or *gdi1*/+ *gdi2* pistils (o). Numbers in brackets indicate displayed/examined embryos. Silhouettes of embryos and division planes are highlighted with dotted lines. Bars = 1 mm (c-g), 10 µm (i-k, m-o).

### Functional loss of *GDI1* and *GDI2* compromises zygotic division and embryonic patterning

To investigate the events leading to embryo lethality of *gdi1 gdi2*, we examined self-fertilized *gdi1*/+ *gdi2* and *gdi1 gdi2*/+ by whole-mount clearing. The ratio of apical to basal cell length at 1-cell stage embryo is a quantitative readout of cell polarization (Mansfield & Briarty, 1991) as it directly reflects the asymmetric positioning of division plane in zygotes (Ueda *et al*., 2017; Wang *et al*., 2021). 1-cell embryos in wild type showed a typical asymmetry, i.e. a small apical cell and a large basal cell (Fig. 2i, 2l). In self-fertilized *gdi1*/+ *gdi2* or *gdi1 gdi2*/+ pistils, 3/4 of 1-cell embryos showed wild-type like apical/basal cell ratio (Fig. 2l). However, around 1/4 of 1-cell embryos were less asymmetric such that the ratio of apical to basal cell length was substantially enlarged (Fig. 2j, 2k).

At later developmental stages, the apical cell divided in a longitudinal orientation to give rise to the embryo whereas the basal cell divides horizontally to give rise to the suspensor in wild type (Fig. 2m), as reported (Lau *et al*., 2012; Dresselhaus & Jurgens, 2021). By contrast, around 1/4 of the 1-cell embryos in *gdi1 gdi2*/+ (Fig. 2n) or *gdi1*/+ *gdi2* (Fig. 2o) showed horizontal instead of longitudinal divisions of the apical cell, leading to subsequent defects in the orientation of division plane, giving rise to a second proembryo-like structure (Fig. 2n-o). These aberrant embryos eventually arrested at the late globular stage (Fig. 2n-o). These results demonstrate that *GDI1* and *GDI2* are crucial for asymmetric zygotic division and for embryonic patterning.

### Functional loss of *GDI1* and *GDI2* compromises cell fate determination after zygotic division

To examine whether cell fate determination after zygotic division was compromised due to functional loss of *GDI1* and *GDI2*, we introduced a marker specific for basal embryo lineage, *pWOX8gΔ:NLS-vYFP3* (Breuninger *et al*., 2008; Ueda *et al*., 2011), in *gdi1/+ gdi2* and examined YFP signals in zygotes and early embryos of the *pWOX8gΔ:NLS-vYFP3 gdi1/+ gdi2* plants. Compared to elongating zygotes in *pWOX8gΔ:NLS-vYFP3* pistils (Fig. 3a), YFP signals were significant reduced in around 25% elongating zygotes (n > 80) within *pWOX8gΔ:NLS-vYFP3 gdi1/+ gdi2* pistils, presumably of the *pWOX8gΔ:NLS-vYFP3 gdi1 gdi2* genotype (Fig. 3b, 3g). From 1-cell embryo stage on, YFP signals were detected in all basal lineage cells in *pWOX8gΔ:NLS-vYFP3* siliques (Fig. 3c, 3e) as reported (Breuninger *et al*., 2008; Ueda *et al*., 2011). By contrast, YFP signals were significantly reduced in around 25% 1-cell embryos (n > 80) within *pWOX8gΔ:NLS-vYFP3 gdi1/+ gdi2* pistils, presumably of the *pWOX8gΔ:NLS-vYFP3 gdi1 gdi2* genotype (Fig. 3d, 3g). At later developmental stages, YFP signals were detected in all basal lineage cells of embryos in *pWOX8gΔ:NLS-vYFP3* (Fig. 3e). Although YFP signals were present in few basal suspensor cells, hypophysis and its directly-subtending suspensor cells in *pWOX8gΔ:NLS-vYFP3 gdi1 gdi2* embryos hardly contained YFP signals (Fig. 3f).

**Figure 3.**
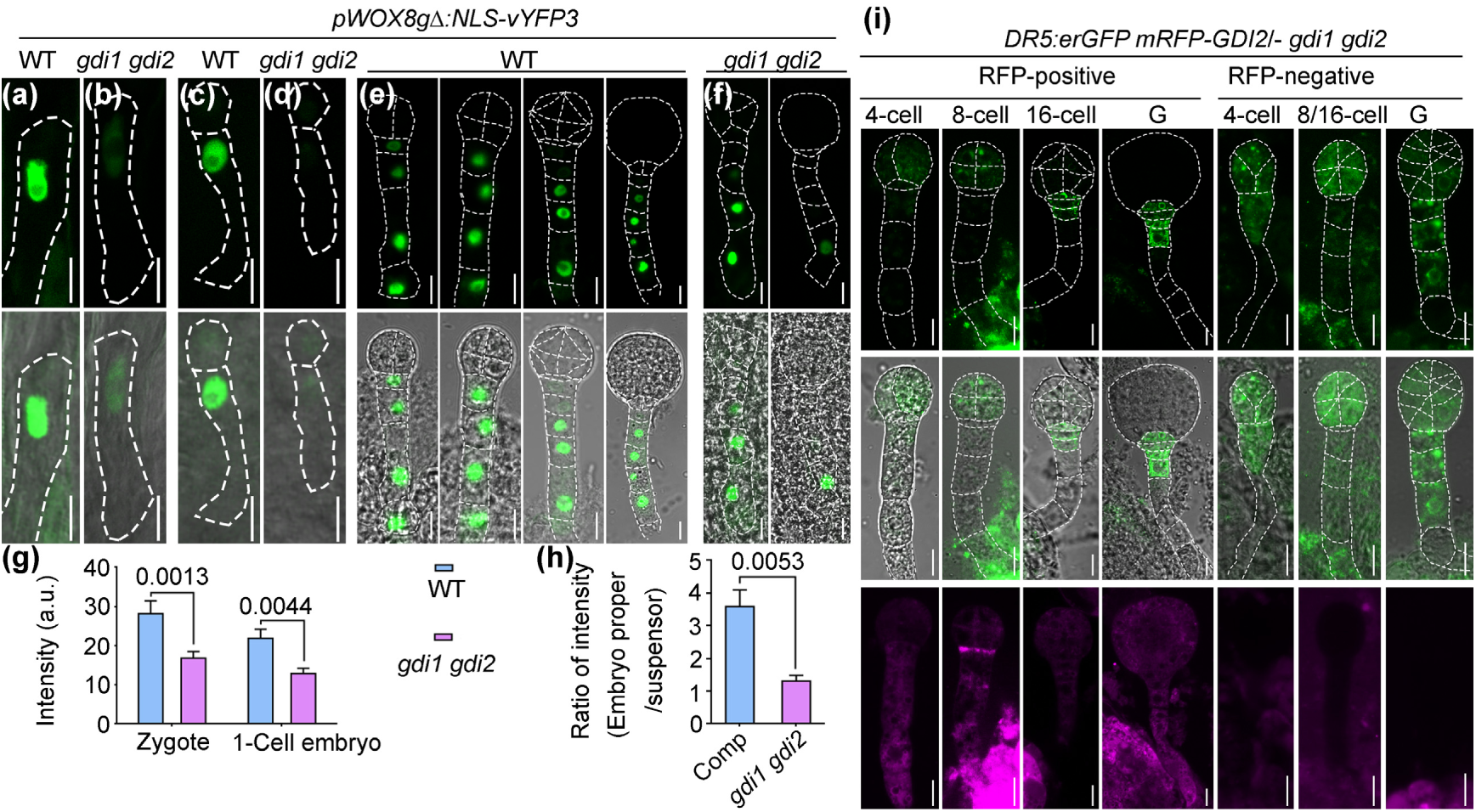
Key factors for early embryogenesis are mis-regulated in *gdi1 gdi2*. (a-f) Representative CLSM images of a zygote (a-b), 1-cell embryo (c-d), or early embryos at various stages (e-f) from the *pWOX8gΔ:NLS-vYFP3* plants (a, c, e) or from the *pWOX8gΔ:NLS-vYFP3 gdi1*/+ *gdi2* plants (b, d, f). Silhouettes of embryos are highlighted with dotted lines. (g) Intensity of NLS-vYFP3 from zygotes or 1-cell embryo of the *pWOX8gΔ:NLS-vYFP3* plants (WT) or the *pWOX8gΔ:NLS-vYFP3 gdi1*/+ *gdi2* plants (*gdi1 gdi2*). a.u. represents arbitrary fluorescence unit. Results are means ± SE (n > 20). P values are shown on top (*t*-test). (h) Ratio of GFP intensity in embryo proper versus in suspensor. Comp indicates RFP-positive embryos whereas *gdi1 gdi2* indicates RFP-negative embryos from the same *DR5:erGFP mRFP-GDI2*/- *gdi1 gdi2* plants. Results are means ± SE (n≥8). P values are shown on top (*t*-test). (i) Representative CLSM images of developing embryos either RFP-positive or RFP-negative from the same *DR5:erGFP mRFP-GDI2*/- *gdi1 gdi2* plants. From top to bottom: the GFP channel images, merges of the GFP and transmission channel images, the RFP channel images. Bars = 10 µm.

Since asymmetric distribution of auxin maximum starting from 1-cell embryo is the other indicator for cell fate determination after the first zygotic division (Friml *et al*., 2003; Rademacher *et al*., 2012; Robert *et al*., 2013; Robert *et al*., 2018), we introduced *DR5:erGFP,* an auxin response reporter (Friml *et al*., 2003), into *mRFP-RabGDI2*/- *gdi1 gdi2* in which zygotes or embryos with RFP signals were of the *mRFP-RabGDI2 gdi1 gdi2* genotype (complemented) whereas those without RFP signals were of the *gdi1 gdi2* genotype. During embryonic patterning, GFP signals were detected in the embryo proper before 16-cells stage embryos (Fig. 3h-i) and in the hypophysis from early globular stages on (Fig. 3i). In contrast to this polar distribution, *gdi1 gdi2* double mutants showed DR5:erGFP signal throughout the embryos (Fig. 3h-i), indicating a compromised auxin polar distribution during embryo development.

Since polar distribution of auxin maximum depends on polar localization of PINs (Benková *et al*., 2003), we introduced *PIN1g:GFP* (Benková *et al*., 2003) into *gdi1/+ gdi2* and selected *PIN1g:GFP gdi1/+ gdi2* plants for analysis. PIN1-GFP signals were detected at newly formed cell membranes in developing embryos of *PIN1g:GFP* (Fig. S6), as reported (Steinmann *et al*., 1999). By contrast, GFP signals were mostly detected in cytoplasmic puncta in *PIN1g:GFP gdi1 gdi2* embryos (Fig. S6). The compromised PM association of PIN1-GFP correlated with altered distribution of auxin maximum (Fig. 3).

Defects both in the expression pattern of *WOX8* and asymmetry of auxin maximum in *gdi1 gdi2* (Fig. 3) demonstrate that functional loss of *GDI1* and *GDI2* compromises cell fate determination immediately after the first zygotic division.

### Mis-targeting of canonical Rab5 and defective vacuolar biogenesis in *gdi1 gdi2* embryos

To determine whether embryo lethality of *gdi1 gdi2* was due to mis-targeting of Rab GTPases, we introduced GFP-ARA7 into the *mRFP-GDI2/- gdi1 gdi2* plants. As a canonical Rab5, ARA7 interacted with RabGDIs (Fig. 1 and Fig. S1). Importantly, functional loss of Vps9a, the GEF for Rab5, resulted in embryo lethality (Goh *et al*., 2007), suggesting the role of Rab5 in embryo development. By comparing the distribution of GFP-ARA7 signals in RFP-positive (*mRFP-GDI2 gdi1 gdi2*, comparable to wild type) and RFP-negative (*gdi1 gdi2*) embryos, we determined that the distribution of ARA7 was affected in *gdi1 gdi2* (Fig. 4). Instead of vesicular distribution in *mRFP-GDI2 gdi1 gdi2* embryos, GFP-ARA7 was distributed as cytoplasmic aggregates in *gdi1 gdi2* embryos (Fig. 4).

**Figure 4.**
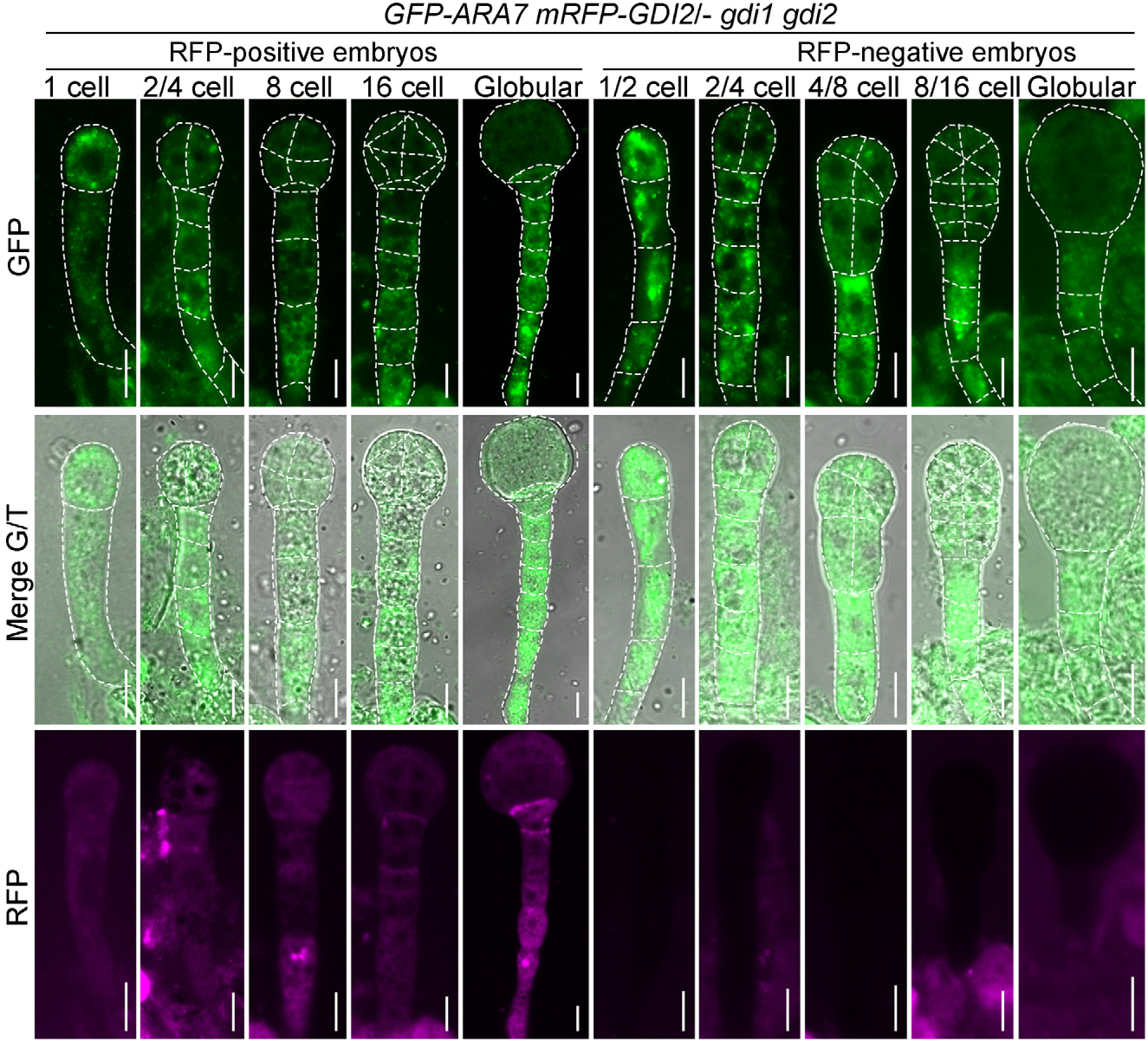
ARA7, the canonical Rab5, is mis-targeted in *gdi1 gdi2*. Representative CLSM images of developing embryos either RFP-positive or RFP-negative from the same *GFP-ARA7 mRFP-GDI2*/- *gdi1 gdi2* plants. From top to bottom: the GFP channel images, merges of the GFP and transmission channel images, the RFP channel images. Silhouettes of embryos are highlighted with dotted lines. Bars = 10 µm.

Because canonical Rab5 is critical for vacuolar trafficking and biogenesis (Minamino & Ueda, 2019) and vacuolar dynamics are critical for zygotic division during embryogenesis (Kimata *et al*., 2019; Matsumoto *et al*., 2021), we wondered whether mis-targeting of ARA7 caused defective vacuolar dynamics in *gdi1 gdi2* embryos, leading to embryo lethality. To test this idea, we examined vacuole morphology by confocal imaging. In bulging zygotes of wild type, vacuoles mostly accumulated at the basal side of bulging zygotes in wild type, enlarged at the basal side to push the nucleus upward in elongating zygotes (Fig. 5a) as reported (Kimata *et al*., 2019). In comparison, bulging zygotes of *gdi1 gdi2* contained randomly distributed vacuoles (Fig. 5b). Large vacuoles could not be detected in elongating *gdi1 gdi2* zygotes (Fig. 5b). Instead, tubular vacuoles extended throughout the elongating *gdi1 gdi2* zygotes with the nucleus randomly positioned (Fig. 5b). In 1-cell embryo of wild type, large vacuoles were present in the basal cell but not apical cell (Fig. 5a). By contrast, the apical cell contained numerous vacuoles whereas vacuoles in the basal cell were fragmented and dispersed throughout the 1-cell *gdi1 gdi2* embryos (Fig. 5b).

**Figure 5.**
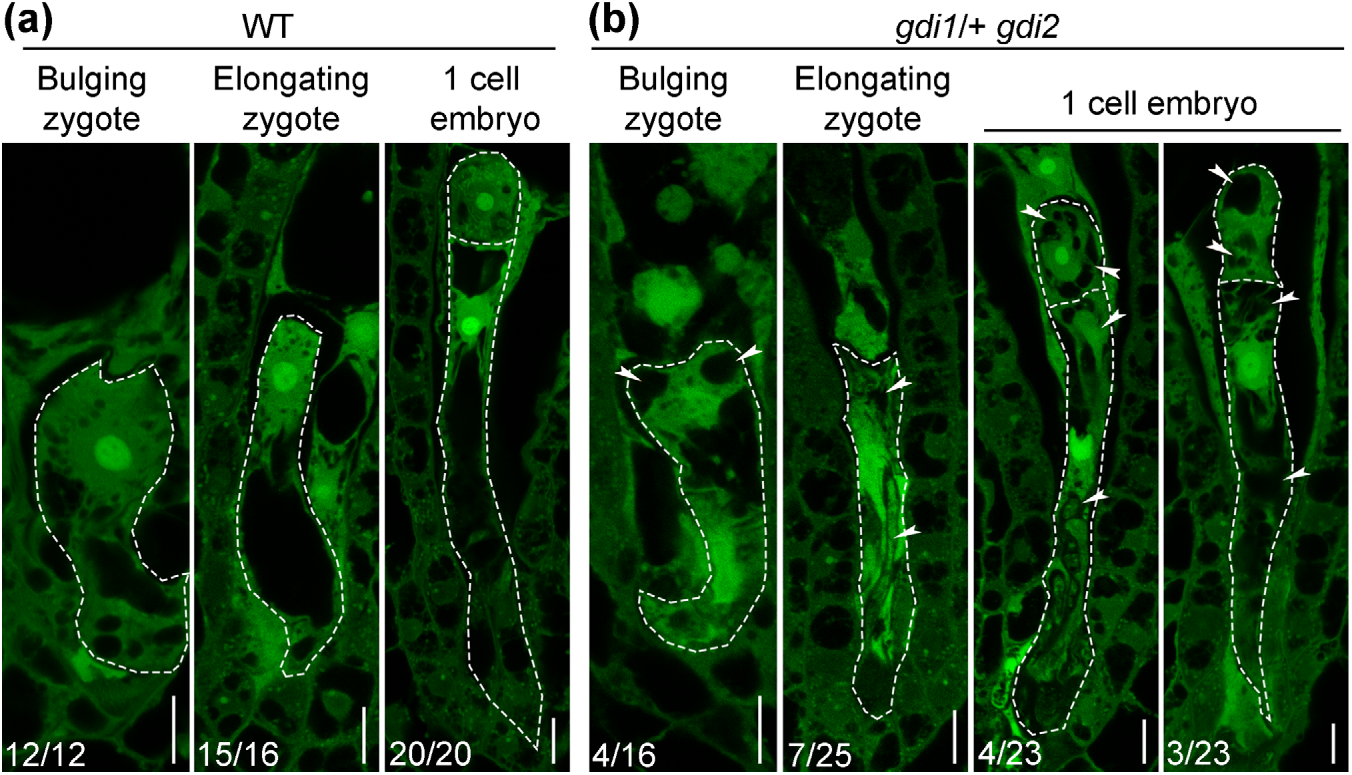
Vacuolar biogenesis and dynamic distribution are compromised in *gdi1 gdi2*. Representative CLSM images of zygotes or 1 cell embryos from wild type (a) or from the *gdi1*/+ *gdi2* plants (b). Silhouettes of zygotes or embryos are highlighted with dotted lines. Numbers at the bottom indicate displayed/examined samples. Arrowheads point at abnormal vacuoles. Bars = 10 µm.

### Genetic interference of canonical Rab5 GTPases mimics the phenotype of *gdi1 gdi2*

To determine whether canonical Rab5 was crucial for GDI-mediated zygotic division and embryonic patterning, we generated *pEC1.1:mRFP-ARA7^S24N^* transgenic plants, in which a dominant negative (DN) form of ARA7 (ARA7^DN^) was driven by a zygote-specific promoter (Xiong *et al*., 2025). We verified the expression of *mRFP-ARA7^DN^* specifically in zygotes based on RFP signals in bulging zygotes (Fig. 6e, 6h), elongating zygotes (Fig. 6f, 6i), as well as in 1-cell embryos (Fig. 6g, 6j). Embryos overexpressing mRFP-ARA7 ^DN^ often showed an enlarged apical/basal cell ratio (Fig. 6b-d), compared to that of wild type (Fig. 6a, 6d), similar to that in *gdi1 gdi2* (Fig. 2). Division patterns of the *pEC1.1:mRFP-ARA7 ^DN^* embryos were abnormal (Fig. 6l-m), similar to but to a less extend than those of *gdi1 gdi2* (Fig. 2).

**Figure 6.**
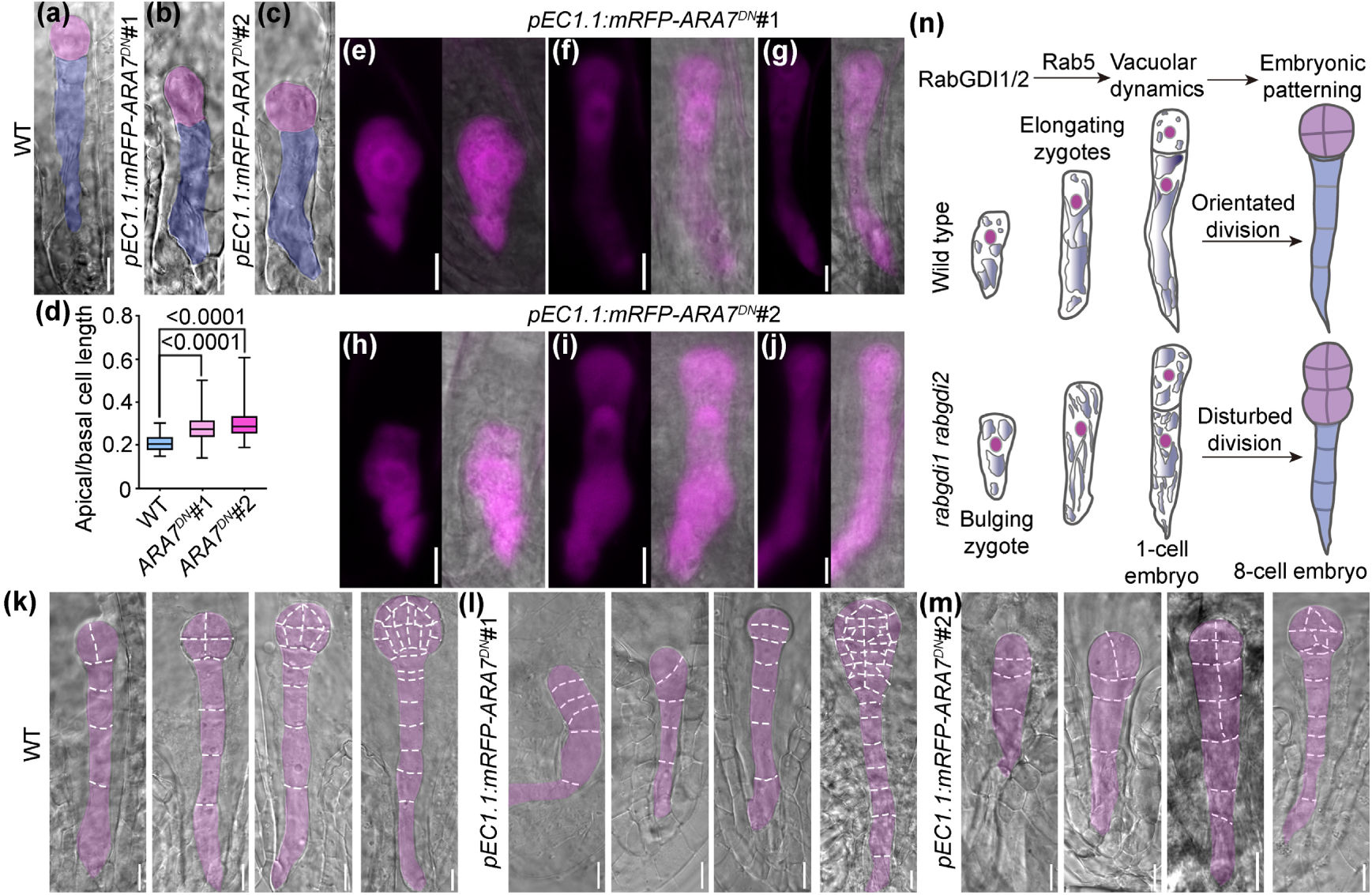
Zygotic expression of a dominant negative Rab5 compromises zygotic division and embryonic patterning. (a-c) DIC images of 1-cell embryo from wild type (a) and two lines of *pEC1.1:mRFP-ARA7^DN^* transgenic plants (b-c). The apical cell and basal cell are pseudo-colored in pink and blue, respectively. (d) Ratio of apical/basal cell length at 1-cell embryos. Results shown are means ± SD (SD, n= 80-132). (e-j) Representative CLSM images of bulging zygotes (e, h), elongating zygotes (f, i), or 1-cell embryos (g, j) from the *pEC1.1:mRFP-ARA7^DN^*#1 (e-g) or the *pEC1.1:mRFP-ARA7^DN^*#2 transgenic plants (h-j). (k-m) Representative DIC images of embryos from 2-cell stages to globular stages from self-fertilized wild-type (k), the *pEC1.1:mRFP-ARA7^DN^*#1 (l), or the *pEC1.1:mRFP-ARA7^DN^*#2 transgenic plants (m). Silhouettes of embryos and division planes are highlighted with dotted lines. (n) A cartoon illustrating the results of this study. Functional loss of *RabGDI1* and *RabGDI2* results in defective vacuolar dynamics, compromises zygotic division and embryonic patterning. Bars = 10 µm.

Because RabGDIs also interacted with the vacuolar trafficking regulator Rab7 and secretory trafficking regulator Rab11 (Fig. 1 and Fig. S1), we wondered whether interfering with the activity of Rab7 or Rab11 by zygotic expression of their DN forms would compromise zygotic division and embryonic patterning. To test this idea, we generated transgenic plants with *pEC1.1:GFP-RabG3f^T22N^* or *pEC1.1:GFP-RabA4b^N130I^*, which expressed the DN form of Rab7 or of Rab11. By CLSM imaging, we verified the expression of GFP-RabG3f^DN^ in bulging zygotes till 1-cell embryos (Fig. S7). However, based on the ratio of apical to basal cell length, there was no significant difference between *pEC1.1:GFP-RabG3f^DN^*transgenic plants and wild type (Fig. S7), suggesting that interfering with the activity of Rab7 in zygotes did not compromise zygotic division. In comparison to the persistent presence of ARA7^DN^ or RabG3f^DN^ in 1-cell embryo (Fig. 6, Fig. S7) and despite the same *pEC1.1* promoter was used, GFP-RabA4b^DN^ signals were undetectable in elongating zygotes or 1-cell embryos (Fig. S7). Also, there was no significant difference between *pEC1.1:GFP-RabA4b^DN^*transgenic plants and wild type in regard to the asymmetric division of zygotes (Fig. S7).

## Discussion

As evolutionarily conserved regulators for dynamic targeting of Rab GTPases, RabGDIs have been identified more than twenty years ago (Zarsky *et al*., 1997; Ueda *et al*., 1998; Ueda *et al*., 2000). Only recently, two maize RabGDI variants were reported to participate in viral resistance through differential binding with a viral protein (Liu *et al*., 2020). Whether and how RabGDIs regulate Rab GTPases in plants remain unclear.

We report that functional loss of two constitutively expressed RabGDI1/2 results in impaired division of zygotes and compromises embryonic patterning (Fig. 6n). Cell fate determination is affected based on the expression of *WOX8* and the polar distribution of auxin (Fig. 3), two major pathways controlling embryogenesis (Friml *et al*., 2003; Haecker *et al*., 2004; Breuninger *et al*., 2008; Rademacher *et al*., 2012; Robert *et al*., 2013; Zhang *et al*., 2017; Robert *et al*., 2018). In addition, vacuolar biogenesis and dynamics that are critical for zygotic division (Kimata *et al*., 2019; Jiang *et al*., 2020) were also compromised in *gdi1 gdi2* (Fig. 5). These results demonstrate a key role of RabGDIs in embryogenesis. RabGDIs specifically interact with canonical Rab GTPases (Fig. 1 and Fig. S1), i.e. Rab GTPases subject to GGT modification (Rutherford & Moore, 2002; Muller & Goody, 2018), but not ARA6 that is a plant-unique Rab5 not modified by GGT (Ueda *et al*., 2001). Functional loss of Rab geranylgeranyl transferase results in embryo lethality in *Arabidopsis* (Rojek *et al*., 2021), supporting a key role of GGT-modified Rab in this process.

Compromised zygotic division and embryonic patterning in *gdi1 gdi2* most likely results from mis-targeting of Rab GTPases, especially canonical Rab5 GTPases. Canonical Rab5 participates in two major vacuolar trafficking routes and is critical for vacuolar biogenesis (Minamino & Ueda, 2019; Hao *et al*., 2023). We show that the canonical Rab5, ARA7, is mis-targeted in zygotes and embryonic cells in *gdi1 gdi2* (Fig. 4). Vacuolar dynamics associated with zygotic polarization and division were compromised in *gdi1 gdi2* (Fig. 5). Consistently, zygotic expression of *ARA7^DN^* results in similar defects at zygotic division and early embryonic patterning to *gdi1 gdi2* (Fig. 6). Indeed, functional loss of *Vps9a*, the major GEF for Rab5, leads to embryo lethality (Goh *et al*., 2007), further supporting a key role of canonical Rab5 GTPases in embryogenesis.

Due to imaging limitations, we were unable to detect the targeting of Rab7 or Rab11 in zygotes or 1-cell embryo at a decent resolution. Rab7 participates in one vacuolar trafficking route (Uemura & Ueda, 2014; Cui *et al*., 2017), interacts with RabGDIs (Fig. 1 and Fig. S1). The representative of Rab7, *RabG3f*, is highly expressed in zygotes (Zhao *et al*., 2019) and decent GFP signals were detected in zygotes or 1-cell embryos of the *pEC1.1:GFP-RabG3f^DN^* transgenic plants (Fig. S7). However, zygotic division and embryonic patterning are not disturbed by RabG3f^DN^ (Fig. S7). One possible explanation is that Rab7 is encoded by multiple members, whose activity would not be easily suppressed by the expression of one Rab7^DN^ form. Alternatively, Rab7-mediated vacuolar trafficking could be compensated for other routes, sufficient to maintain vacuolar dynamics during zygotic division.

Zygotic expression of the DN-formed RabA4b did not interfere with zygotic division (Fig. S7) despite that RabA4b is a key regulator for secretory trafficking (Nielsen *et al*., 2008) and a member of Rab11 highly expressed in zygotes (Zhao *et al*., 2019). However, we demonstrate that PINs are mis-targeted in *gdi1 gdi2* (Fig. S6), which is consistent with disrupted auxin distribution after zygotic division in *gdi1 gdi2* (Fig. 3). Polar PM targeting of PINs depends on secretory trafficking (Geldner *et al*., 2001; Geldner *et al*., 2003; Hille *et al*., 2018), which is not only regulated by Arf GTPases (Geldner *et al*., 2001; Geldner *et al*., 2003) but also by Rab GTPases (Nielsen *et al*., 2008). Thus, functional loss of RabGDIs most likely interferes with secretory trafficking. The un-disturbed zygotic division in the *pEC1.1:GFP-RabA4b^DN^* transgenic plants is likely due to the rapid turnover of RabA4b^DN^ as judged by the absence of GFP signals in elongating zygotes or 1-cell embryos of the *pEC1.1:GFP-RabA4b^DN^* transgenic plants (Fig. S7). Alternatively, RabA4b-mediated post-Golgi secretion represents only one of multiple secretory pathways existing for zygotic division and embryonic patterning.

## Author contributions

Y. Z. and F. X. conceived and supervised the project; G.-M. Y. and Y.-N. W. performed the experiments with the assistance of S.-S. D., S. L.; W. W., L. J., and Z.-z. L. assisted with imaging, Y. Z., F. X., and G.-M. Y. designed the experiments and analyzed the data; Y. Z., G.-M. Y., and F. X. wrote the article with contributions from all the authors. All authors read and approved of the manuscript.

## Acknowledgement

We thank Prof. Ruixi Li for *pDONR221-P1P4-TOPO*, *pDONR221-P3P2-TOPO*, *pBiFCt-2in1-NN* and *pBiFCt-2in1-NC* vectors. This work has been supported by grants from Natural Science Foundation of China (32461160286, 32270805, 32470807, 32400290), by Shandong Provincial Natural Science Foundation (ZR2024MC093), by the Research Grants Council of Hong Kong (N_CUHK482/24), and by Taishan Scholar Project of Shandong Province of China (tsqn202408133). The authors declare no competing financial interests.

