## Supplemental figures for "Arabidopsis RabGDIs are essential for the asymmetric division of zygotes and embryonic patterning"

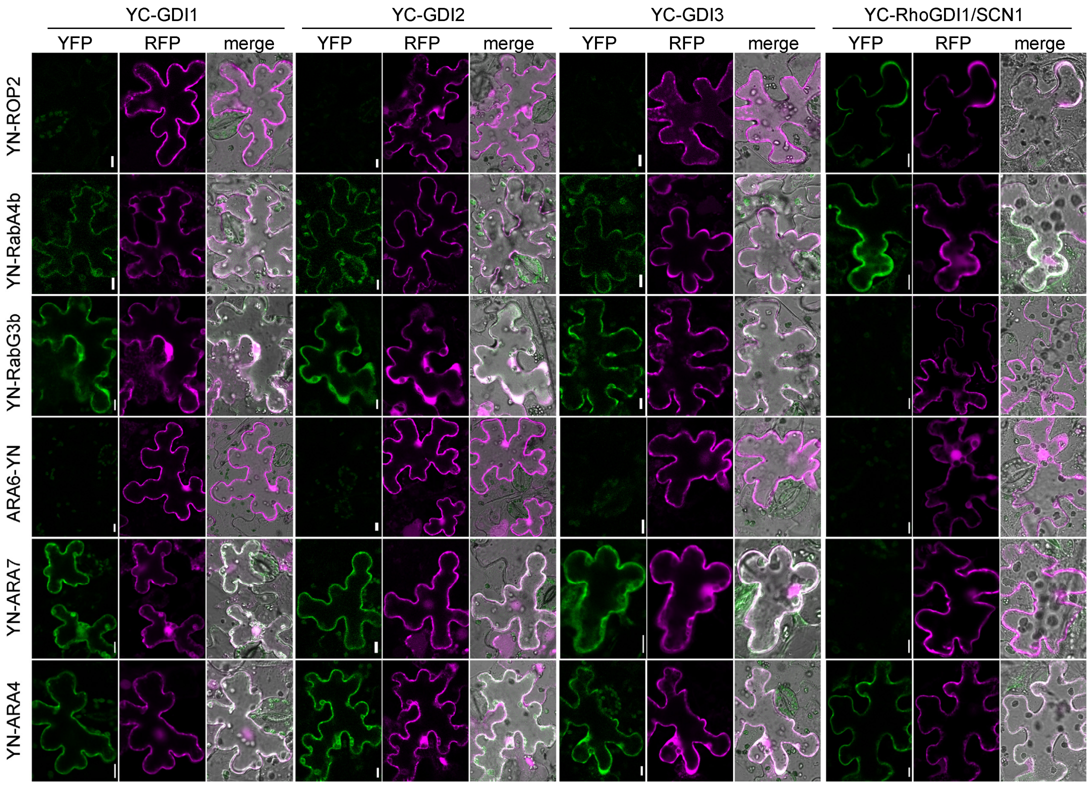


**Supplemental Figure 1. Interaction between RabGDIs and Rab GTPases.**
Representative BiFC assays to examine the interaction between RabGDIs and related Rab GTPases. RhoGDI1/SCN1 was used as the control for RabGDIs whereas ROP2 was used as the control for Rab GTPases. Positive signals were indicated by signals in the YFP channels. RFP signals indicate co-expressed mRFP. Bars = 10 µm.

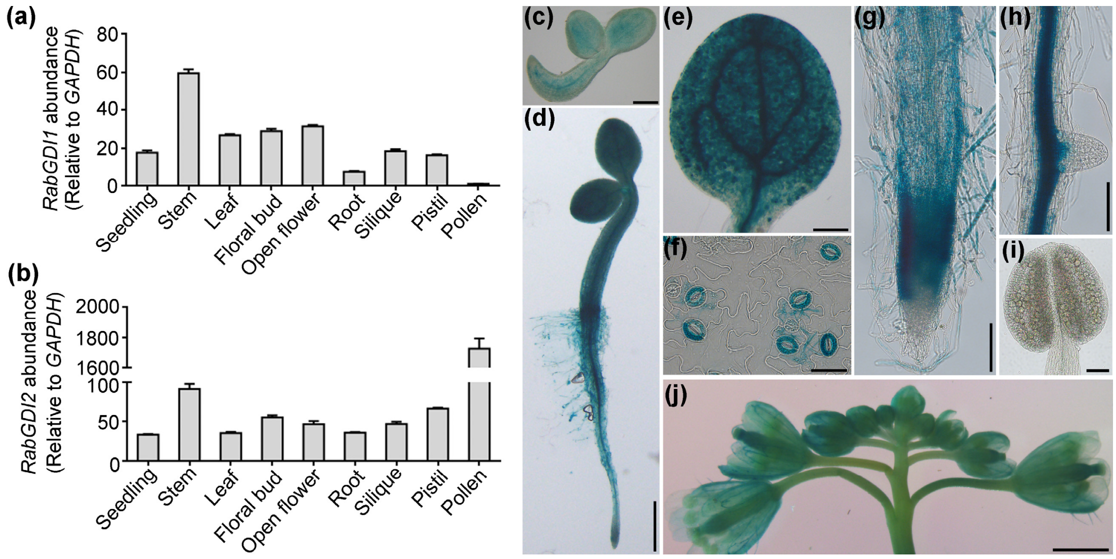


**Supplemental Figure 2. *RabGDI1* and *RabGDI2* are constitutively expressed.**
(a-b) Quantitative real-time PCRs of *GDI1* (a) or *GDI2* (b) among different tissues. Results shown are means ± standard errors (SE, n=3). Experiments were repeated three times with similar results. (c-j) Representative histochemical GUS staining of a seedling at 24 hrs after germination (HAG, c) or 48 HAG (d), a leaf (e) and leaf epidermal peel (f), a primary root (g), a lateral root (h), a mature anther (i), or an inflorescence (j) from *GDI1g:GUS* plants. Bars = 200 µm for (c, e), 500 µm for (d), 20 µm for (f), 50 µm for (g-i), 1 mm for (j).

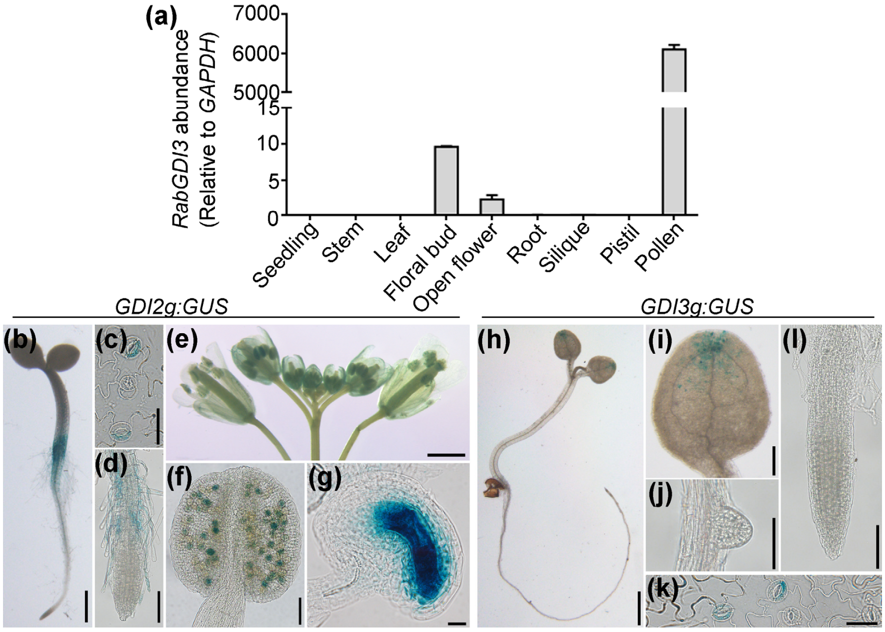


**Supplemental Figure 3. Expression of *RabGDI2* and *RabGDI3* in other tissues than pollen.**

(a) Quantitative real-time PCRs of *GDI3* among different tissues. Results shown are means ± standard errors (SE, n=3). Experiments were repeated three times with similar results. (b-g) Representative histochemical GUS staining of a seedling at 48 hrs after germination (b), an epidermal peel (c), a primary root (d), an inflorescence (e), a mature anther (f), or a mature ovule (g) from *GDI2g:GUS* plants. (h-l) Representative histochemical GUS staining of a seedling at 4 days after germination (DAG, h), a cotyledon (i), a lateral root (j), an epidermal peel (k), or a primary root (l) from *GDI3g:GUS* plants. Bars = 1 mm for (e, h), 500 µm for (b), 200 µm for (i), 50 µm for (d, f, j, l), 20 µm for (c, g, k).

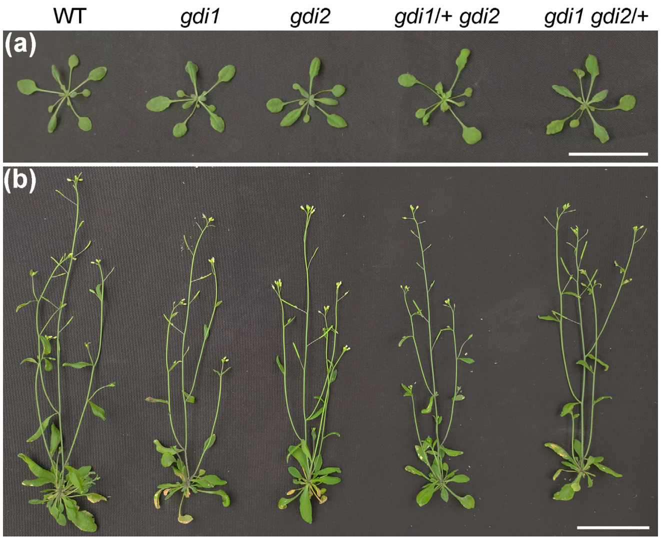


**Supplemental Figure 4. Plant growth is not compromised in *gdi1*/+ *gdi2* or *gdi1 gdi2*/+.**

(a-b) Representative plants of various genotypes at 2 WAG (a) or 5 WAG (b). Bars = 5 cm.

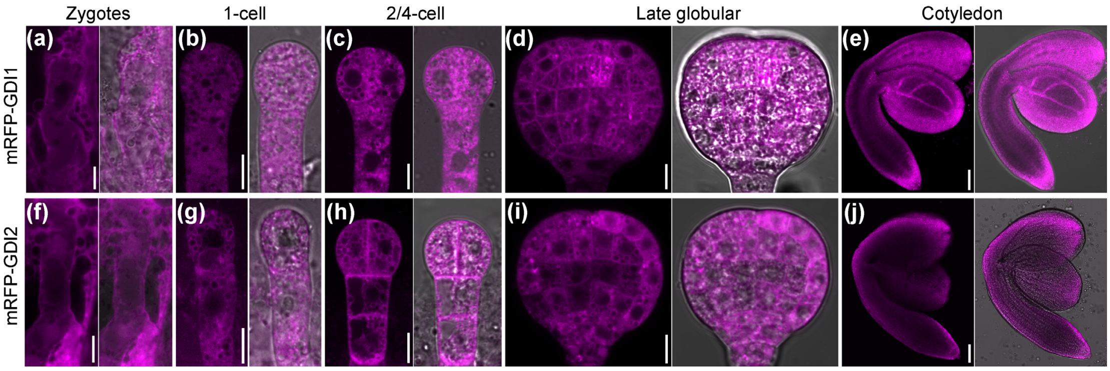


**Supplemental Figure 5. Expression of mRFP-GDI1 or mRFP-GDI2 fully rescues the embryo lethality of *gdi1 gdi2*.**

(a-j) Representative CLSM images of an elongating zygote (a, f), a 1-cell stage embryo (b, g), a 2 to 4-cell stage embryo (c, h), an embryo at late globular stage (d, i), or an embryo at the cotyledon stage (e, j) from the *proGDI1:mRFP-GDI1 gdi1 gdi2* plants (a-e) or from the *proGDI2:mRFP-GDI2 gdi1 gdi2* plants (f-j). Bars = 10 μm (a-d, f-i), 50 μm (e, j).

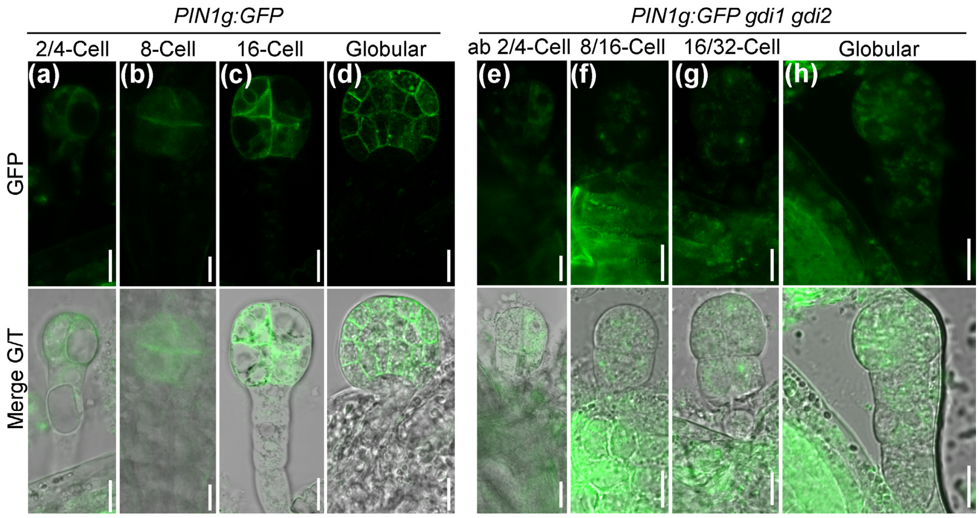


**Supplemental Figure 6. PM localization of PIN1 at embryo proper cells is compromised by *RabGDI1* and *RabGDI2* loss-of-function.**

(a-h) Representative CLSM images showing PIN1-GFP in *PIN1g:GFP* (a-d) or in *PIN1g:GFP gdi1 gdi2* embryos (e-h) at various stages. Ab, about. Top: the GFP channel; bottom: merges of the GFP and transmission channels. Bars = 10 µm.

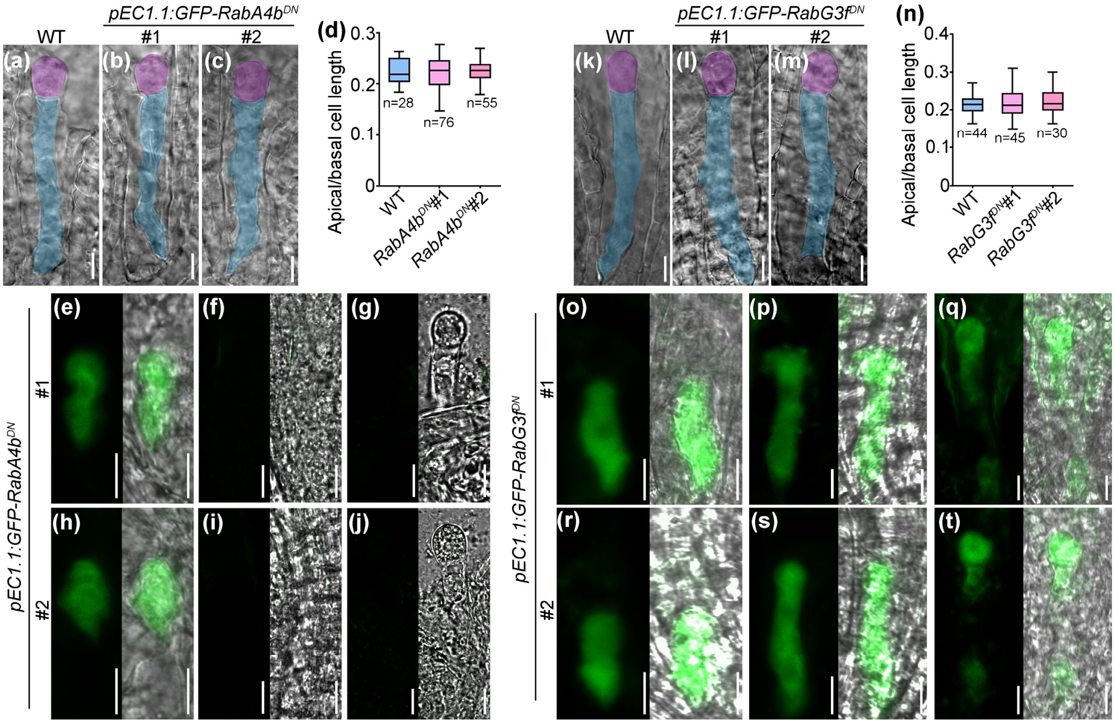


**Supplemental Figure 7. Zygotic expression of Rab11$^{DN}$ or Rab7$^{DN}$ does not compromise zygotic division.**

(a-c) Representative DIC images of 1-cell embryo from wild type (a) or two lines of *pEC1.1:GFP-RabA4b$^{DN}$* transgenic plants (b-c). (d) Ratio of apical/basal cell length at 1-cell embryos. Results shown are means ± SD (SD, n= 28-76). (e-j) CLSM images of a bulging zygote (e, h), an elongating zygote (f, i), or 1-cell embryo (g, j) from two lines of *pEC1.1:GFP-RabA4b$^{DN}$* transgenic plants. Right are merges of the GFP/transmission channels. (k-m) Representative DIC images of 1-cell embryo from wild type (k) or two lines of *pEC1.1:GFP-RabG3f$^{DN}$* transgenic plants (l-m). (n) Ratio of apical/basal cell length at 1-cell embryos. Results shown are means ± SD (SD, n= 30-45). (o-t) CLSM images of a bulging zygote (o, r), an elongating zygote (p, s), or 1-cell embryo (q, t) from two lines of *pEC1.1:GFP-RabG3f$^{DN}$* transgenic plants. Right are merges of the GFP/transmission channels. For (a-c, k-m), the apical cell and basal cell are pseudo-colored in pink and blue, respectively. All groups are not significantly different from each other for (d, n) (OneWay ANOVA, Tukey's multiple comparisons test, P>0.05). Bars = 10 μm.

**Supplemental Table 1. Oligos used in this study.**

| Application | | No. | Sequence 5’-3’ |
|---|---|---|---|
| Cloning for expression pattern | *GDI1p* | ZP9867 | ATGTGTGGATTTTGAGTTAGTCATCGTTCTCCATCCACTCAG |
| | | ZP9868 | ACCCGGGGATCGATCCTGCAGTTGCTGATCCTCCAAAGAGAA |
| | *GDI2p* | ZP9871 | ATGTGTGGATTTTGAGTTAGGAGAGGCTGAGAAAAACCCTGT |
| | | ZP9872 | ACCCGGGGATCGATCCTGCGGCTGATTTTCGTAGAGCAAACG |
| | *GDI3p* | ZP9875 | ATGTGTGGATTTTGAGTTAGTCAGCTGCAACTCTCGATCG |
| | | ZP9876 | ACCCGGGGATCGATCCTGCTATTGTAATCGATCTCTATCTCCGTTATTG |
| | *GDI1g* | ZP9869 | CGGGAATTGCGTACCGAGCTATGGATGAAGAGTACGAAGTCATAGTT |
| | | ZP9870 | CGATCGGGGAAATTCGAGCTTCATTCCTCCTCTGCAGCAC |
| | *GDI2g* | ZP9873 | CGGGAATTGCGTACCGAGCTATGGATGAAGAGTACGAGGTTATTGTTC |
| | | ZP9874 | CGATCGGGGAAATTCGAGCTTCATTCCTCTGCAGCACTTGC |
| | *GDI3g* | ZP9877 | CTCGGGAATTGCTACCGAGCTCATGGATGAAGAGTATGATGTGATTGTC |
| | | ZP9878 | CGATCGGGGAAATTCGAGCTTCAATTCTCTGCAGCAGCAC |
| qRT-PCR | *GDI1* | ZP9500 | CTCATTTAGGTTCTAGCAGAGACTA |
| | | ZP9501 | TAAAAGACAAGTACTTGGTCACATC |
| | *GDI2* | ZP9502 | TAAGTTTATGATGGGAAATGGC |
| | | ZP9503 | CTTTTGAACCTTTCCTTTGAC |
| | *GDI3* | ZP9504 | GCGAGAAAGTTCTTTATCTATGT |
| | | ZP9505 | TTGTGTCATCTTCAAGTCCATA |
| | *GAPDH* | ZP687 | TGAAATCAAAAAGGCTATCAAGG |
| | | ZP688 | CATCATCCTCGGTGTATCCAA |
| CRISPR/Cas9 | *GDI1* and *GDI2* | ZP8219 | ATATATGGTCTCGATTGTGGACAGGAATGACTACTAGTT |
| | | ZP8220 | TGTGGACAGGAATGACTACTAGTTTTAGAGCTAGAAATAGC |
| | | ZP8221 | AACTAGTAGTCGTTTCTATCCACAATCTCTTAGTCGACTCTAC |
| | | ZP8222 | ATTATTGGTCTCGAAACTAGTAGTCGTTTCTATCCACAA |
| Sequencing | *GDI1g* | P3705 | CCATGCTTGATCTGGAGATTTTCAA |
| | | PZ234 | GGAATAGGTACGCATGATGAGATCG |
| | *GDI2g* | ZP9249 | TTGCTTCTTCTAATAGACTTCTTC |
| | | ZP9250 | TCATCAACAATTTAAGAGTTATCAG |
| | | PZ235 | ATATAAAAGAAAACAACACAAGGTT |
| Cloning for complement | *GDI1* CDS | ZP10584 | GCCACTCCACCGGCGCCGGAATGGATGAAGAGTACGAAGTCATAGTTCT |
| | | ZP10585 | TGAACGATCGGGGAAATTCGTCATTCCTCCTCTGCAGCACTAG |
| | *GDI2* CDS | ZP10586 | GCCACTCCACCGGCGCCGGAATGGATGAAGAGTACGAGGTTATTGTTC |
| | | ZP10587 | TGAACGATCGGGGAAATTCGTCATTCCTCTGCAGCACTTGC |
| | *GDI1p* | P4564 | ATGTGTGGATTTTGAGTTAGTTGTCGACCAGCCAGGATTAAA |

| | | | |
|---|---|---|---|
| | | | CGAGGTAGAG |
| | | P4565 | CTCGGAGGAGGCCATACTAGTAGTTGCTGATCCTCCAAAGAGAA |
| | *GDI1g* | P4568 | CACTCCACCGGCGCCGGATCCATGGATGAAGAGTACGAAGTCATAGTT |
| | | P4569 | ACGATCGGGGAAATTCGGATCCTCATTCCTCCTCTGCAGCAC |
| | *GDI2p* | P4566 | ATGTGTGGATTTTGAGTTAGTTGTCGACCCTTGTCAAGTCTGTGACTTTC |
| | | P4567 | CTCGGAGGAGGCCATACTAGTGGCTGATTTTCGTAGAGCAAAC |
| | *GDI2g* | P4570 | GCCGCCACTCCACCGGCGCCGGATCCATGGATGAAGAGTACGAGGTTATTGTTC |
| | | P4571 | ACGATCGGGGAAATTCGGATCCTCATTCCTCTGCAGCACTTGC |
| Cloning for entry vectors | *ROP2* CDS | ZP492 | CACCATGGCGTCAAGGTTTATAAAGT |
| | | ZP493 | TCACAAGAACGCGCAAC |
| | *ARA6* CDS | ZP1390 | CACCATGGGATGTGCTTCTTCTCTT |
| | | ZP2039 | TGACGAAGGAGCAGGACGAGGTAG |
| | *RabG3b* CDS | ZP927 | CACCATGTCGACGCGAAGACGAAC |
| | | ZP928 | TCAGCAAGCACAACCTCCTC |
| | *ARA7* CDS | PZ1 | CACCATGGCTGCAGCTGGAAACAA |
| | | ZP8378 | CTAAGCACAACAAGATGAGCTCACTG |
| | *RabA4b* CDS | ZP3762 | CACCATGGCCGGAGGAGGC |
| | | ZP3763 | TCAAGAAGAAGTACAACAAGTGCTG |
| | *ARA4* CDS | ZP9338 | TTTAAGAAGGAGCCCTTCACCATGTCAGACGACGACGAGAGAGG |
| | | ZP9340 | GGGTCGGCGCGCCCACCCTTTTACCTCGAACAGCAAGAGAATGTCTTTG |
| | *GDI1* CDS | ZP8397 | CACCATGGATGAAGAGTACGAAGTCATAGTTCT |
| | | ZP9497 | TCATTCCTCCTCTGCAGCACTAG |
| | *GDI2* CDS | ZP8399 | CACCATGGATGAAGAGTACGAGGTTATTGTTC |
| | | ZP9498 | TCATTCCTCTGCAGCACTTGC |
| | *GDI3* CDS | ZP8401 | CACCATGGATGAAGAGTATGATGTGATTGTCC |
| | | ZP9499 | TCAATTCTCTGCAGCAGCACTTG |
| Cloning for BiFC vectors | *ROP2* CDS | P210 | GGGGACAACTTTGTATAATAAAGTTGGAATGGCGTCAAGGTTTATAAAGTGT |
| | | P211 | GGGGACCACTTTGTACAAGAAAGCTGGGTTCACAAGAACGCGCAACG |
| | *ARA6* CDS | P1133 | GGGGACAACTTTGTATAATAAAGTTGGAATGGGATGTGCTTCTTCTCTT |
| | | P1134 | GGGGACCACTTTGTACAAGAAAGCTGGGTGTGACGAAGGAGCAGGACG |
| | *RabG3b* | P212 | GGGGACAACTTTGTATAATAAAGTTGGAATGTCGACGCGAAGA |

| | | | |
|---|---|---|---|
| | CDS | | CGAA |
| | | P213 | GGGGACCACTTTGTACAAGAAAGCTGGGTTCAGCAAGCACAACCTCCT |
| | *ARA7* CDS | ZP11522 | GGGGACAACTTTGTATAATAAAGTTGGAATGGCTGCAGCTGGAAACAAGAGCATTAA |
| | | ZP11523 | GGGGACCACTTTGTACAAGAAAGCTGGGTCTAAGCACAACAAGATGAGCTCACT |
| | *RabA4b* CDS | ZP28Y | GGGGACAACTTTGTATAATAAAGTTGGAATGGCCGGAGGAGGC |
| | | ZP29Y | GGGGACCACTTTGTACAAGAAAGCTGGGTTCAAGAAGAAGTACAACAAGTGCT |
| | *ARA4* CDS | P208 | GGGGACAACTTTGTATAATAAAGTTGGAATGTCAGACGACGACGAGA |
| | | P209 | GGGGACCACTTTGTACAAGAAAGCTGGGTTTACCTCGAACAGCAAGAGAATG |
| | *GDI1* CDS | ZP11516 | GGGGACAAGTTTGTACAAAAAAGCAGGCTTAATGGATGAAGAGTACGAAGTCA |
| | | P216 | GGGGACAACTTTGTATAGAAAAGTTGGGTTCATTCCTCCTCTGCAGCACTAGC |
| | *GDI2* CDS | ZP11518 | GGGGACAAGTTTGTACAAAAAAGCAGGCTTAATGGATGAAGAGTACGAGGTTATTG |
| | | P217 | GGGGACAACTTTGTATAGAAAAGTTGGGTTCATTCCTCTGCAGCACTTGCGGCA |
| | *GDI3* CDS | ZP11520 | GGGGACAAGTTTGTACAAAAAAGCAGGCTTAATGGATGAAGAGTATGATGTGATTGTCC |
| | | P218 | GGGGACAACTTTGTATAGAAAAGTTGGGTTCAATTCTCTGCAGCAGCACTTGC |
| | *SCN1* CDS | ZP78Y | GGGGACAAGTTTGTACAAAAAAGCAGGCTTAATGTCTTTGGTATCTGGAGCC |
| | | ZP79Y | GGGGACAACTTTGTATAGAAAAGTTGGGTTCAAAGCGCAGGCCATT |
| Cloning for *EC1.1p:RFP-ARA7$^{DN}$* | *EC1.1p* | P5020 | ATGTGTGGATTTTGAGTTAGTTGTCGACCTATCATGAATTAGCTCTACTAAATCT |
| | | P5021 | CTCGGAGGAGGCCATACTAGTTTCTCAACAGATTGATAAGGTCG |
| | *ARA7$^{S24N}$ CDS* | J717 | GATGTTGGTGCTGGAAAAAACAGTCTTGTGTTACGGTTTGT |
| | | J718 | TTTTCCAGCACCAACATCTCCAAGCAACACCTGAA |
| | | P5026 | GCCGCCACTCCACCGGCGCCGGATCCATGGCTGCAGCTGGAAACA |
| | | P5027 | ACGATCGGGGAAATTCGGATCCCTAAGCACAACAAGATGAGCTCAC |
| Cloning for *EC1.1p:GFP-RabG3f$^{DN}$* | *EC1.1p* | P5023 | AACTGCAGCTATCATGAATTAGCTCTACTAAATCT |
| | | P5024 | CGGGATCCTTCTCAACAGATTGATAAGGTCG |
| | *RabG3f$^{T22N}$ CDS* | ZP2545 | CACCATGCCGTCCCGTAGACGTAC |
| | | ZP2546 | TTAGCATTCACACCCTGTAGACCT |

| | | | |
|---|---|---|---|
| | | ZP2739 | GGTGGGAAAAAACTCTTTGATG |
| | | ZP2740 | CCGCTATCACCGAGGATG |
| | | PZ201 | GTACATGGTAGATCTTGAGCTCATGCCGTCCCGTAGACGTAC |
| | | PZ202 | CGATCGGGGAAATTCGAGCTCTTAGCATTCACACCCTGTAGACCT |
| Cloning for *pEC1.1:GFP-RabA4b$^{DN}$* | *EC1.1p* | PZ231 | TGATCCAAGCTCAAGCTAAGCTTCTATCATGAATTAGCTCTACTAAATCT |
| | | PZ233 | TCGCCCTTGCTCACCATACTAGTTTCTCAACAGATTGATAAGGTCG |
| | *RabA4b $^{N130I}$ CDS* | ZP1089 | CATCCTTATTGGAATCAAGTCTGATCTAG |
| | | ZP1090 | ATGACAATGTTCTTATCAGCGTG |
