## Supplemental Table for "Arabidopsis RabGDIs are essential for the asymmetric division of zygotes and embryonic patterning"

**Table 1. Gametophyte transmission of *rabgdi1 gdi2* and segregation ratio of heterozygous mutants.**

| **Female × Male** | **genotype** | **Expected ratio** | **Observed ratio** |
| --- | --- | --- | --- |
| *gdi1^+/-^* *gdi2* × wild type | *gdi1^+/-^*: *^+/+^* (*gdi2^+/-^*) | 1: 1 | 39: 31 |
| wild type × *gdi1^+/-^* *gdi2* | *gdi1^+/-^*: *^+/+^* (*gdi2^+/-^*) | 1: 1 | 37: 25 |
| *gdi1* *gdi2^+/-^* × wild type | *gdi2^+/-^*: *^+/+^* (*gdi1^+/-^*) | 1: 1 | 27: 26 |
| wild type × *gdi1* *gdi2^+/-^* | *gdi2^+/-^*: *^+/+^* (*gdi1^+/-^*) | 1: 1 | 28: 27 |
| *gdi1^+/-^* × *gdi1^+/-^* | *gdi1^+/+^*: *^+/-^*: *^-/-^* | 1: 2: 1 | 16: 31: 13 |
| *gdi2^+/-^* × *gdi2^+/-^* | *gdi2^+/+^*: *^+/-^*: *^-/-^* | 1: 2: 1 | 15: 26: 13 |
| *gdi1^+/-^ gdi2* × *gdi1^+/-^ gdi2* | *gdi1^+/+^*: *^+/-^*: *^-/-^* (*gdi2*) | 1: 2: 1 | 40: 62: 0 ^a^ |
| *gdi1 gdi2^+/-^* × *gdi1* *gdi2^+/-^* | *gdi2^+/+^*: *^+/-^*: *^-/-^*(*gdi1*) | 1: 2: 1 | 51: 93: 0 ^a^ |

^a^ Significantly different from the segregation ratio 1: 2: 1 (χ^2^, p<0.01).
